# *In vivo* single-cell profiling of lncRNAs during Ebola virus infection

**DOI:** 10.1101/2022.01.12.476002

**Authors:** Luisa Santus, Raquel García-Pérez, Maria Sopena-Rios, Aaron E Lin, Gordon C Adams, Kayla G Barnes, Katherine J Siddle, Shirlee Wohl, Ferran Reverter, John L Rinn, Richard S Bennett, Lisa E Hensley, Pardis C Sabeti, Marta Melé

**Affiliations:** Life Sciences Department, Barcelona Supercomputing Center, Barcelona, Catalonia 08034, Spain; FAS Center for Systems Biology, Department of Organismic and Evolutionary Biology, Harvard University, Cambridge, MA 02138, USA; Broad Institute of MIT and Harvard, Cambridge, MA 02142, USA; Harvard Program in Virology, Harvard Medical School, Boston, MA 02115, USA; Department of Organismic and Evolutionary Biology, Harvard University, Cambridge, MA, USA; Integrated Research Facility, Division of Clinical Research, National Institute of Allergy and Infectious Diseases, National Institutes of Health, Frederick, MD 21702, USA; Department of Epidemiology, Johns Hopkins Bloomberg School of Public Health, Baltimore, United States; Department of Genetics, Microbiology and Statistics University of Barcelona, Barcelona, Spain; Department of Immunology and Infectious Diseases, Harvard T.H. Chan School of Public Health, Harvard University, Boston, MA 02115, USA; Howard Hughes Medical Institute, Chevy Chase, MD 20815, USA; Department of Biochemistry, University of Colorado Boulder, Boulder 80303, USA

**Keywords:** Long non-coding RNAs, Ebola virus, Single-cell, Immune response, *De novo* transcriptome assembly

## Abstract

Long non-coding RNAs (lncRNAs) are pivotal mediators of systemic immune response to viral infection, yet most studies concerning their expression and functions upon immune stimulation are limited to *in vitro* bulk cell populations. This strongly constrains our understanding of how lncRNA expression varies at single-cell resolution, and how their cell-type specific immune regulatory roles may differ compared to protein-coding genes. Here, we perform the first in-depth characterization of lncRNA expression variation at single-cell resolution during Ebola virus (EBOV) infection *in vivo*. Using bulk RNA-sequencing from 119 samples and 12 tissue types, we significantly expand the current macaque lncRNA annotation. We then profile lncRNA expression variation in immune circulating single-cells during EBOV infection and find that lncRNAs’ expression in fewer cells is a major differentiating factor from their protein-coding gene counterparts. Upon EBOV infection, lncRNAs present dynamic and mostly cell-type specific changes in their expression profiles especially in monocytes, the main cell type targeted by EBOV. Such changes are associated with gene regulatory modules related to important innate immune responses such as interferon response and purine metabolism. Within infected cells, several lncRNAs have positively and negatively correlated expression with viral load, suggesting that expression of some of these lncRNAs might be directly hijacked by EBOV to attack host cells. This study provides novel insights into the roles that lncRNAs play in the host response to acute viral infection and paves the way for future lncRNA studies at single-cell resolution.

## Introduction

EBOV is one of the most lethal pathogens to humans and it is infamously notorious for its high infectiousness and severe case fatality rates (Malvy et al. 2019; Jacob et al. 2020). In the past, EBOV caused alarming outbreaks and up to the present day, it represents a major global health threat (Ilunga Kalenga et al. 2019). Previously, bulk tissue transcriptomic analyses improved our understanding of EBOV’s evoked host immune response (Caballero et al. 2016; Jain et al. 2020). Now, emerging single-cell RNA sequencing (scRNA-Seq) technologies are refining our understanding of the systemic immune response mounted upon viral infections (Jain et al. 2020; Kazer et al. 2020; Kotliar et al. 2020) by allowing the dissection of gene expression dynamics in multiple cell populations simultaneously. More importantly, in the case of organisms infected with a virus, scRNA-Seq can identify and profile infected cells separately from uninfected bystander cells, and thus, distinguish the host cellular transcriptional response triggered by viral replication versus the inflammatory cytokine milieu. Indeed, recent work on circulating immune cell types upon EBOV infection *in vivo* revealed that although host defenses are characterized by strong activation of innate immunity, within infected cells, EBOV evades cell’s defenses by suppressing antiviral gene expression allowing for uncontrolled viral replication (Messaoudi et al. 2015; Kotliar et al. 2020). However, previous studies have focused on the host protein-coding gene response and have systematically ignored the role that non-coding genes such as lncRNAs may play in the host response to EBOV infection. This is mostly due to poor lncRNA annotations in non-human primates, the main species of EBOV research.

Although understudied in the context of EBOV infection, lncRNAs are crucial host immune response regulators (Heward and Lindsay 2014; Marina R. Hadjicharalambous 2019). LncRNAs regulate the maturation and development of lymphoid and myeloid cells which are pivotal cell lineages in the immune response. Specifically, lncRNAs mediate hematopoietic stem cells’ differentiation (Luo et al. 2015), quiescence maintenance (Venkatraman et al. 2013) and survival (Kotzin et al. 2016). In addition, lncRNAs are involved in the first line of defense of the innate immune response as they mediate pathogen-induced monocytes and macrophages activation and the subsequent release of inflammatory factors such as cytokine and chemokines (Marina R. Hadjicharalambous 2019; Mariotti et al. 2019; Cui et al. 2019). Finally, they have been shown to participate in antiviral responses, such as interferon signalling (Suarez et al. 2020) and their transcriptional profiles have been shown to dynamically change upon infection of several viral pathogens (Fortes and Morris 2016).

The mechanism of action of lncRNAs, together with many intrinsic properties, distinguish lncRNAs from protein coding genes. LncRNAs often regulate gene expression by acting as signaling molecules (Nicholas W. Mathy 2017; Wang and Chang 2011; Pandey et al. 2008), decoys (Kallen et al. 2013), molecular guides (Grote et al. 2013) or through scaffolding (Yang et al. 2014). In addition, despite lncRNAs share similar biogenesis with protein-coding genes (Quinn and Chang 2016; Derrien et al. 2012), they are distinguishable by a variety of features, such as lower expression levels (Cabili et al. 2011; Igor Ulitsky 2013; Derrien et al. 2012), higher tissue specificity (Derrien et al. 2012; Cabili et al. 2011, 2015; Hezroni et al. 2015) and lower splicing efficiency (Tilgner et al. 2012; Melé et al. 2017). However, most of these observations arise from bulk tissue analyses; therefore, whether their expression kinetics is driven by overall low expression levels across many cells or by high expression levels in specific cell populations remains unclear. This lack of knowledge at single cell resolution hampers our understanding of how lncRNAs function and whether their regulation and response upon infection is intrinsically different from that of protein coding genes.

In this work, we significantly expand lncRNA macaque annotation to study lncRNA expression dynamics in circulating immune single-cells infected with EBOV *in vivo*. The goal of this work is to identify lncRNAs that may be playing crucial roles in the context of EBOV host response and that may be manipulated by EBOV to induce viral replication.

## Results

### *De novo* annotation largely expands the rhesus macaque non-coding transcriptome

Bulk and single-cell transcriptomic studies in rhesus macaque reported widespread host gene expression changes upon EBOV infection (Siragam et al. 2018; Nakayama and Saijo 2013; Kotliar et al. 2020). However, most lncRNAs have been systematically excluded in such studies due to incomplete annotations, especially in rhesus monkey, where the number of annotated lncRNAs reaches only 28% of that in human (Supplemental Fig. S1A). To improve the current lncRNA annotation, we generated RNA-sequencing for 12 tissues of healthy and EBOV infected macaques. We additionally combined this data with publicly available blood RNA-sequencing of healthy and EBOV infected macaques (Cross et al. 2018), adding up to a total of 119 samples and almost 4 billion reads (Supplemental Table S1). To identify novel lncRNAs, we implemented a computational pipeline that performs *de novo* transcriptome assembly, extensive quality controls, and non-coding transcript selection based on concordance between three different tools (Figure 1B, Supplemental Fig. S1B). Our approach had high accuracy (82%) and specificity (86%) when predicting Ensembl annotated macaque lncRNAs (Supplemental Fig. S1C). In total, we discovered 3,979 novel lncRNA genes (5,299 transcripts) (Figure 1C), of which 3,191 (80%) were intergenic and 788 (20%) were antisense. Consistent with previous studies (Bryzghalov et al. 2019), we identified a human lncRNA ortholog for 528 lncRNAs (14%) (Supplemental Fig. S1D). As expected, novel and annotated lncRNA transcripts were shorter and had fewer and longer exons compared to protein-coding genes (Mann-Whitney U test, all P-values < 2.2 x10^−16^) (Derrien et al. 2012; Uszczynska-Ratajczak et al. 2018) (Figure 1D-F). However, novel lncRNAs were shorter than annotated lncRNAs in macaque (Mann-Whitney U test, all P-values < 2.2 x 10^−16^). To discard the possibility that our pipeline detected incomplete transcripts, we selected novel lncRNAs that had a human ortholog and compared transcript lengths between species. Novel lncRNAs were not shorter than their human orthologs (Paired Wilcoxon signed-rank test, P-value > 0.05) (Supplemental Fig. S1E), suggesting that our novel lncRNA annotation captures full-length transcripts. As expected, both annotated and novel lncRNAs are expressed in fewer tissues and show lower expression levels than protein-coding genes (Chen et al. 2018; Cabili et al. 2011, 2015; Hezroni et al. 2015) (Figure 1G-H). These observations hold when we analyze intergenic and antisense lncRNAs separately (Supplemental Fig. S2A-E). In summary, our novel lncRNAs resemble lncRNA reference annotations and almost double current rhesus macaque reference lncRNA annotation.

**Figure 1.**
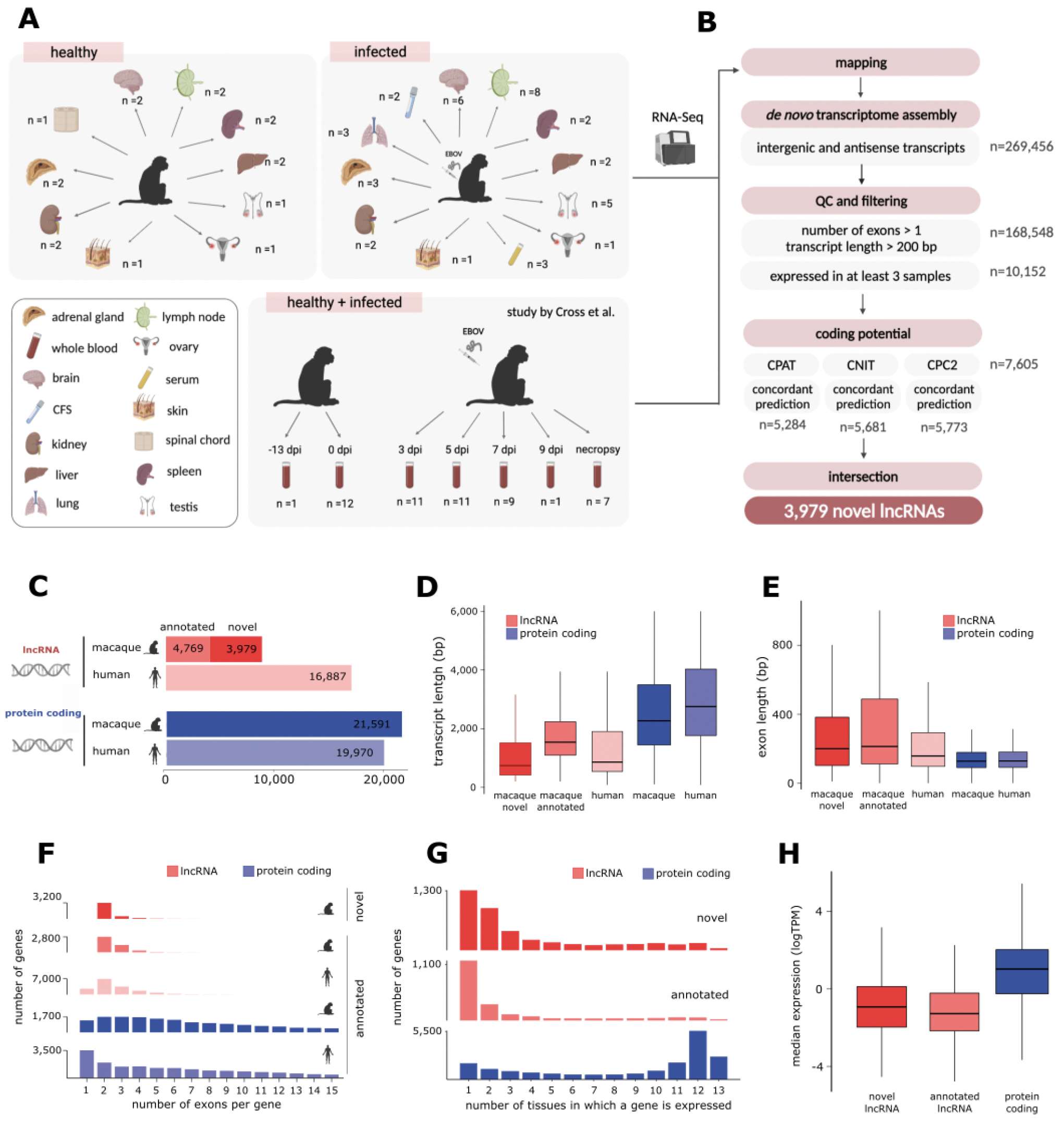
Novel lncRNAs resemble annotated lncRNAs and significantly expand the current macaque lncRNA annotation. **(A)** Tissue samples used for *de novo* transcriptome assembly. **(B)** LncRNA discovery pipeline. The number of lncRNA genes that pass each filtering step are indicated. **(C)** Number of novel and annotated lncRNA and protein-coding genes in the macaque and human annotation (Ensembl release 100). **(D)** Transcript length, **(E)** exon length**, (F)** number of exons per transcript novel and annotated macaque and human lncRNA (red) and protein-coding genes (blue). (**G**) Number of tissues in which macaque genes are expressed and (**H**) median expression levels of macaque novel and annotated lncRNA (red) and protein-coding genes (blue).

### LncRNAs are systematically expressed in fewer cells compared to protein-coding genes

LncRNAs are more lowly expressed, more tissue-specific, and often have a more time-dependent expression in comparison to protein-coding genes. However, these observations have mostly been made in bulk tissue studies (Cabili et al. 2011; Derrien et al. 2012; Hezroni et al. 2015; Cabili et al. 2015; Melé et al. 2017), and thus, whether this signal arises from lncRNAs being lowly expressed within individual cells or from their expression being restricted to only a few cells remains elusive (Katerina AB Gawronski 2017). To address this, we used single-cell transcriptomics data from macaque’s peripheral blood mononuclear cells (PBMCs) from Kotliar et al. (Kotliar et al. 2020; Bennett et al. 2020). We selected 38,067 cells through rigorous quality control (see Methods) and classified them into 4 major cell-types: monocytes, neutrophils, B cells and T cells (Figure 2A, Supplemental Fig. S3A). We found that the main feature distinguishing lncRNAs from protein-coding genes, rather than their median single-cell expression level (Figure 2B), is that lncRNAs are expressed in fewer cells (Mann-Whitney U test, P-value = 0.017 and P-value < 2.2 x10^−16^ respectively) (Figure 2C). See figure 2D-E for an example where a lncRNA (Figure 2D) and a protein-coding gene (Figure 2E) are expressed at comparable median expression levels but differ in the number of cells in which they are expressed. When we inspected cell types separately (Supplemental Fig. S3B-C), we also found that the biggest difference between the two gene classes is driven by the fraction of cells in which they are expressed.

**Figure 2.**
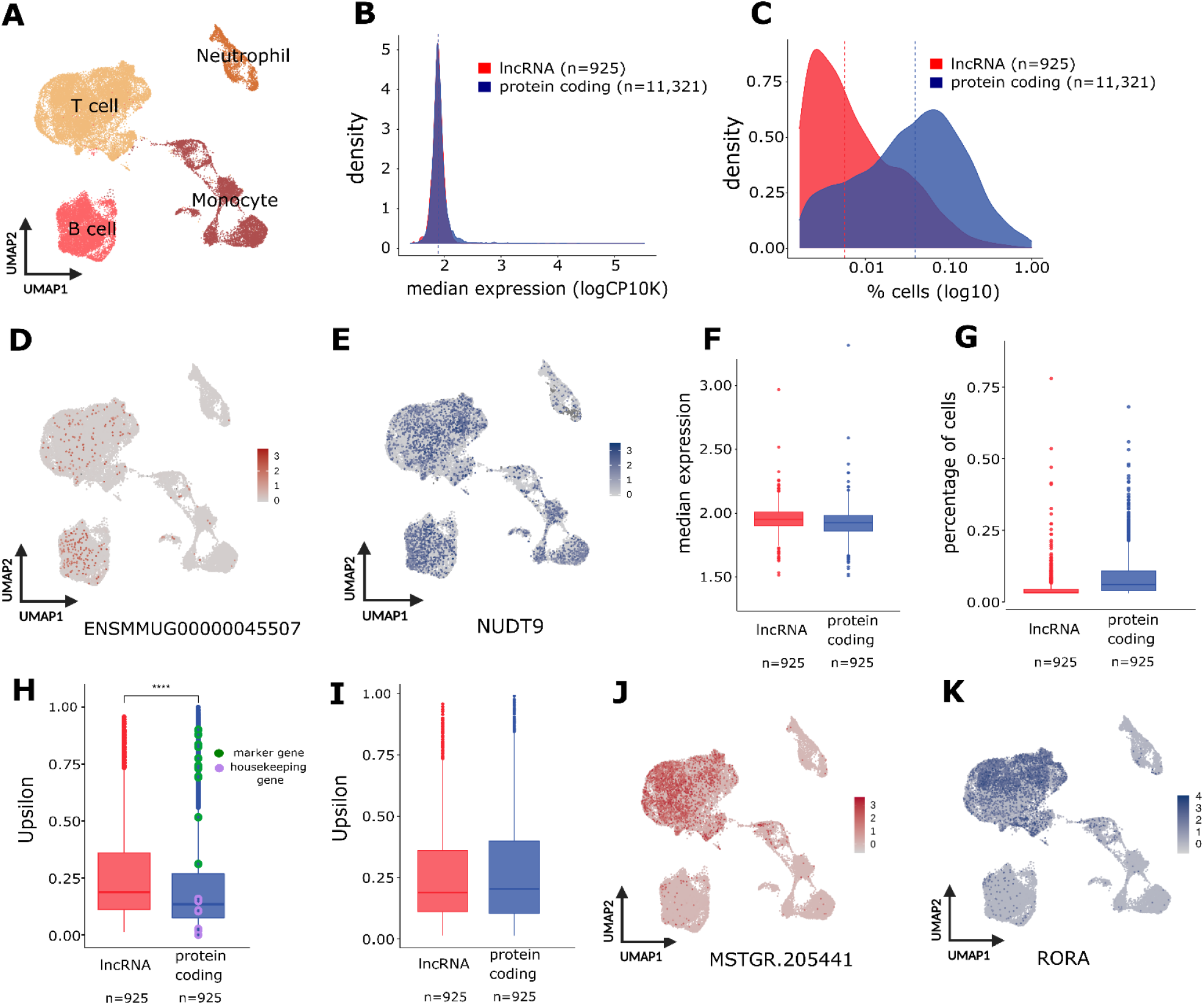
Expression patterns of lncRNAs and protein-coding genes at single-cell resolution. **(A**) UMAP embedding of 38,067 cells. Cell-types are indicated by the different colours. **(B)** Median expression values (logCP10K) and **(C)** percentage of cells (log10) at which lncRNA (red) and protein-coding genes (blue) are expressed. **(D)** UMAP embedding showing the expression levels of a lncRNA (ENSMMUG00000045507) and **(E)** a protein coding gene (NUDT9), with the same median expression levels but expressed in a different number of cells. **(F)** Median expression levels of lncRNAs (red) and protein-coding genes (blue) when matched by the percentage of cells in which they are expressed. **(G)** Percentage of cells in which lncRNAs (red) and protein-coding genes (blue) are expressed when matched by median expression levels. **(H)** Boxplot comparing the cell-type specificity scores of lncRNAs (red) and protein-coding genes (blue). Cell-type marker genes are highlighted in green, housekeeping genes in purple. **(I)** Upsilon specificity score of lncRNAs and protein-coding genes when matched by the percentage of cells in which they are expressed. **(J)** UMAP embedding showing cell-type-specific of lncRNA MSTRG.205441 (Upsilon = 0.9) and **(K)** protein-coding gene RORA (Upsilon =0.89), which were matched by the number of cells in which they are expressed.

The proportion of cells in which each gene is expressed and its expression levels are tightly correlated (Supplemental Fig. S3D). To control for this, we tested whether lncRNA expression levels were lower than those of protein-coding genes when expressed in a comparable number of cells. We found no evidence of protein-coding genes being expressed at higher levels than lncRNAs when matched by the proportion of cells in which they are expressed (one-side Wilcoxon signed-rank test, P-value > 0.05) (Figure 2F). Conversely, lncRNAs were expressed in fewer cells compared to protein-coding genes when controlling for the median expression levels of the two gene classes (one-side Wilcoxon signed-rank test, P-value < 2.2 x 10^−16^) (Figure 2G). To assess whether our observations are consistent regardless of single-cell technology, quality of the gene annotation or infection status, we replicated these analyses using healthy human PBMCs generated with a different platform (10X Genomics) and observed the same trends (Supplemental Fig. S3G-I).

LncRNAs are more tissue-specific than protein-coding genes (Cabili et al. 2011; Igor Ulitsky 2013). However, this pattern could be explained by a greater cell-type specificity of lncRNAs or by lncRNA expression being restricted to fewer cells independent of cell type. To address this, we used the established metric Tau (Kryuchkova-Mostacci and Robinson-Rechavi 2017) and also designed a new metric, Upsilon (see Methods), to estimate the cell-type specificity per gene. Whereas Tau relies mostly on differences in expression levels (Kryuchkova-Mostacci and Robinson-Rechavi 2017), Upsilon relies exclusively on the proportion of cells in which each gene is expressed. Both Tau and Upsilon could correctly classify genes as highly, intermediate or low cell-type specific in simulated scenarios (Methods and Supplemental Fig. S4). As expected, cell marker genes had high cell-type specificity scores whereas housekeeping genes had low specificity scores for both metrics. Importantly, Upsilon provided a better separation for intermediate cell-type specificity stages as well as from marker to housekeeping genes (Figure 2H, Supplemental Fig. S4A, S5A). Novel lncRNAs showed similar cell-type specificity scores compared to annotated lncRNAs (Mann-Whitney test, P-value > 0.05). Notably, lncRNAs had higher cell-type specificity than protein-coding genes (Mann-Whitney U test, P-value < 2 x10^−10^) (Figure 2H, Supplemental Fig. S5A), but when matched by the number of cells in which they were expressed, they had comparable cell-type specificity scores (Wilcoxon signed-rank test, P-value > 0.05) (Figure 2I, Supplemental Fig. S5C). To assess whether these observations were independent of species, annotation, infection status, or sequencing platform, we analyzed healthy human PBMC single-cell data generated with the 10X Genomics platform and our results were consistent (Supplemental Fig. S5E-F).

Overall, our results suggest that in circulating immune cells, lower expression levels of lncRNA compared to protein-coding genes reported in bulk studies could be driven by lncRNAs being expressed in fewer cells and that lncRNAs are as cell-type specific as protein-coding genes when expressed in the same number of cells.

### Cell-type dynamic regulation of lncRNAs upon EBOV infection

LncRNAs play crucial roles in the host response to viral infections (Ginn et al.; Kesheh et al.; Fortes and Morris 2016; Liu and Ding 2017). However, previous studies mostly focused on specific cell lines or bulk tissues. Cell lines analyses are reductive of *in vivo* biology, whereas bulk tissue data is not ideal as it averages gene expression and detects with higher difficulty genes expressed only in few cells, such as lncRNAs. To overcome these limitations, we leveraged the previously introduced scRNA-Seq dataset. To identify lncRNAs that may play important cell-type specific immune regulatory roles upon viral infection we tested for differential gene expression, comparing each stage of the infection (early, middle, late) to baseline in each cell type separately (monocytes, T and B cells)(see Methods). We detected a total of 110 differentially expressed (DE) lncRNAs in at least one cell type (FDR <0.05, fold change >30%) (Figure 3A-E)(Supplemental Table S2), the majority of which (73 lncRNA, 66%) were novel, underscoring the importance of refining the annotation of lncRNAs for model organisms such as rhesus monkeys. The largest number of DE lncRNA was in monocytes (74 DE genes)(Figure 3C, Supplemental Fig. S6A), consistent with monocytes being the main EBOV target (Geisbert et al. 2003b, 2003a). Most lncRNAs (99 lncRNAs, ~90%) were differentially expressed in exclusively one cell type (Figure 4F); this is a slightly larger proportion than the one observed for protein-coding genes (Fisher’s exact test, OR = 2.77, P-value = 0.00028; see Methods). However, when matched by the proportion of cells in which they are expressed, the two gene classes had comparable proportions of cell-type specifically differentially expressed genes (Fisher’s exact test; OR = 1.1, P-value = 0.5). Also, most lncRNAs (88 lncRNAs, ~80%) were differentially expressed in only one stage of the infection (Figure 4G) which is a significantly larger 7 proportion to that of protein-coding genes, (Fisher’s exact test, OR = 3.21, P-value = 1.009 x10^−7^; see Methods). However, when comparing genes matched by the number of cells in which they are expressed, the differences between lncRNAs and protein coding genes in stage-specific differential expression were reduced (Fisher’s exact test; OR = 2.2, P-value = 0.005).

**Figure 3.**
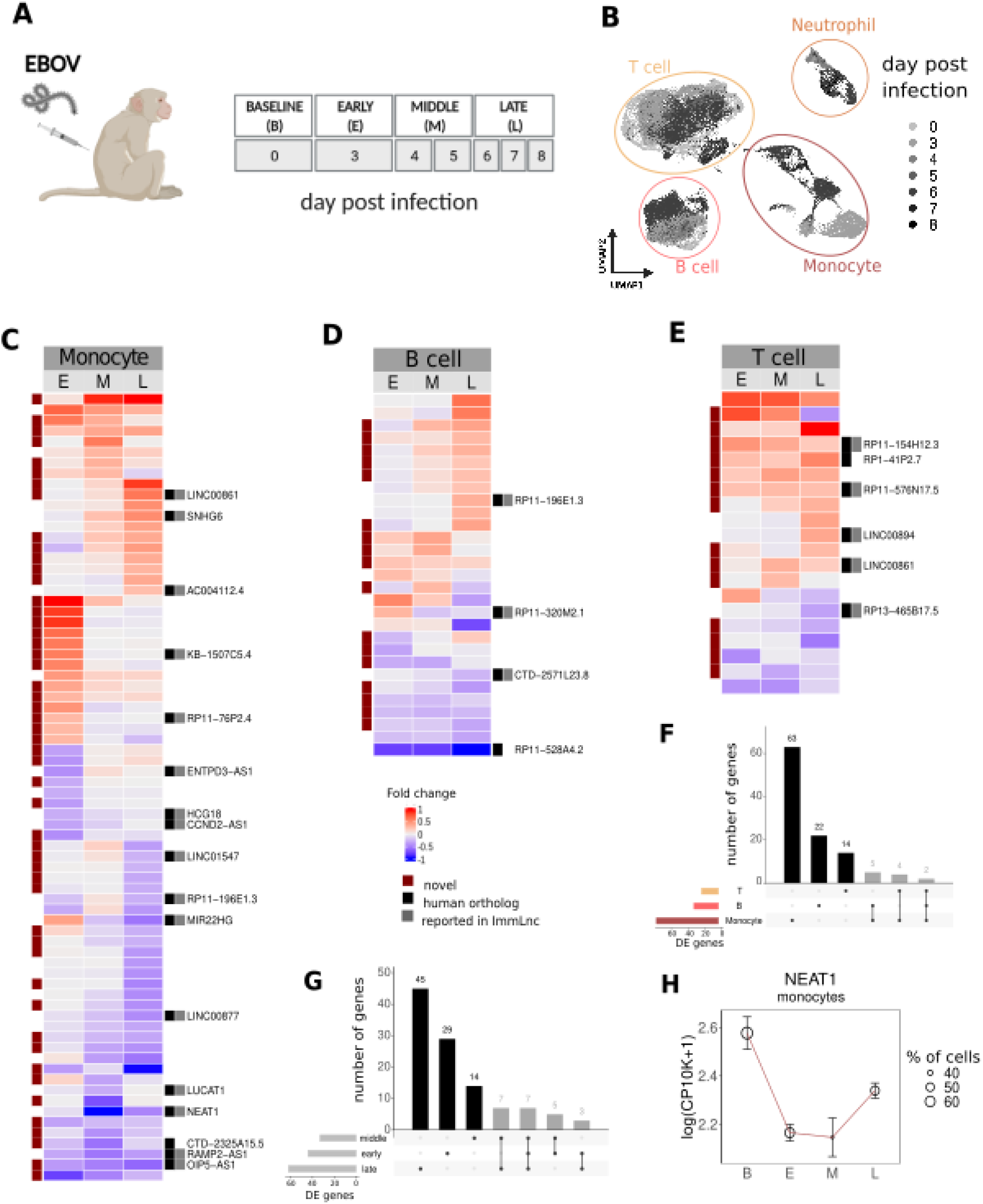
LncRNA expression changes upon EBOV infection are cell-type specific. **(A)** Schematic overview for the *in vivo* experiment design. **(B)** UMAP embedding of 38,067 cells from the *in vivo* dataset, coloured by sampling day relative to infection day. Heatmap coloured by fold changes (log2) of differentially expressed genes in early (E), middle (M) and late (L) stages of the infection in Monocytes (**C**), T cells (**D**) and B cells (**E**). Upset plot showing the overlap of lncRNAs differentially expressed across cell types (**F**) and disease progression stages (**G**). **(H)** Expression patterns of the NEAT1 at different stages of infection in monocytes, B and T cells. Marker: mean; error bars: 95% confidence interval. Dot’s sizes represent the percentage of cells in which the gene is expressed in each cell type.

**Figure 4.**
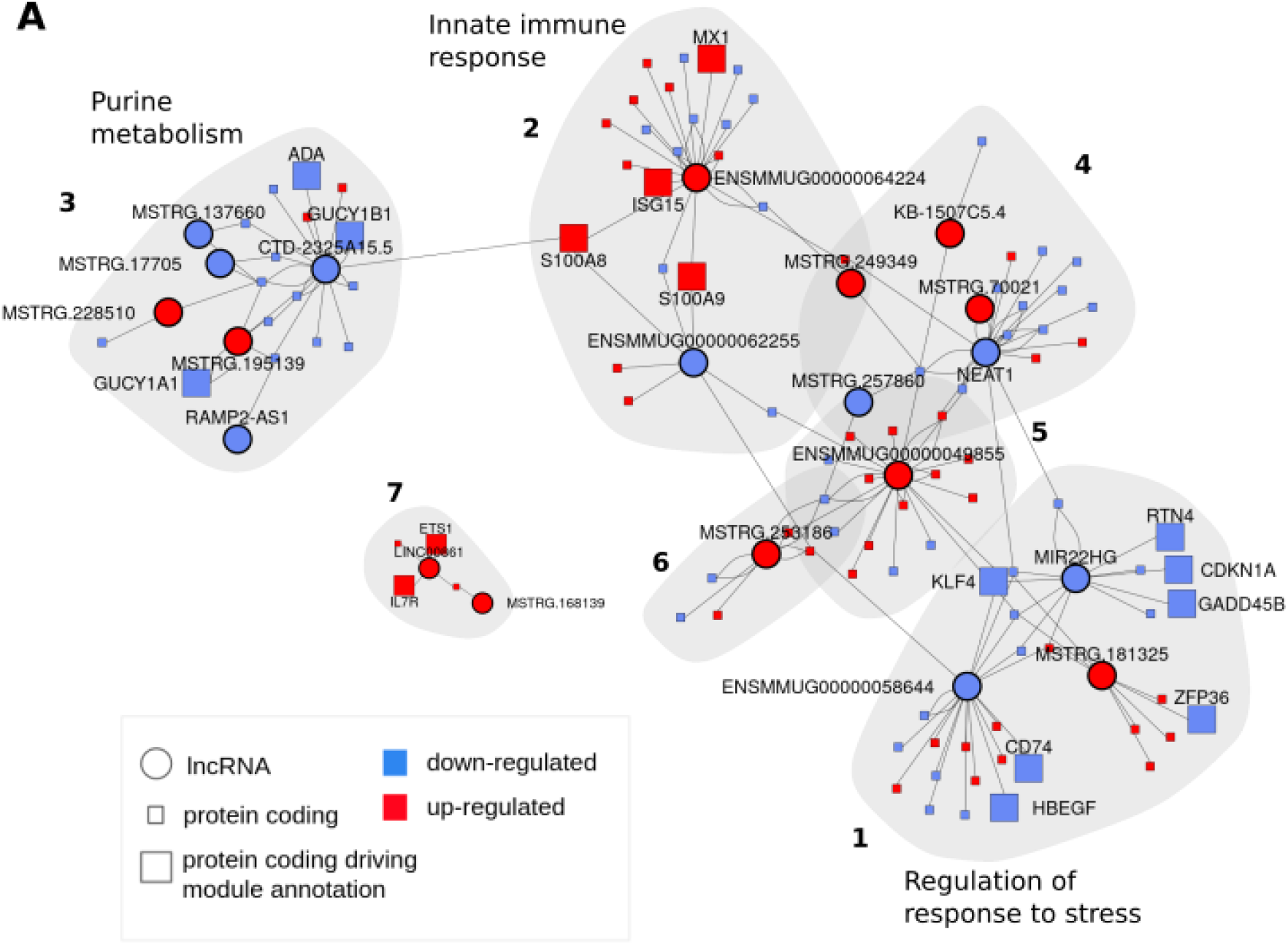
Co-expression network of lncRNAs and protein-coding genes upon EBOV infection in monocytes. **(A)** Regulatory network of DE lncRNAs (circles) and DE protein-coding genes (squares). Vertices’ colours represent whether a gene is up- or down-regulated upon EBOV infection. Protein-coding genes driving the functional enrichment of the module and lncRNAs are highlighted in bigger sizes.

Twenty-five DE lncRNA had a human ortholog, and, reassuringly, twenty-two of those had been previously reported to change expression during immune response in humans (Li et al. 2020b) (Figure 3C-E, Supplemental Fig. S6E). Consistent with previous studies regarding immune response upon infection, LINC00877 (Razooky et al. 2017; Policard et al. 2021) was downregulated, and SNHG6 upregulated (Zhao et al. 2016; Waickman et al. 2019). Interestingly, the most transcriptionally repressed lncRNA was the nuclear enriched abundant transcript 1 (NEAT1) (Figure 3C, H). NEAT1 is a well-studied lncRNA known to play important anti-viral roles. In most studies, however, NEAT1 is upregulated upon viral infections (Prinz et al. 2019) and downregulation has only been described in dengue and Crimean Congo hemorrhagic fever (Pandey et al. 2017; Bayyurt et al. 2021). Our results suggest that NEAT1 depletion may be specific to severe hemorrhagic fevers and it is the first time that such downregulation is shown to occur specifically in monocytes.

Overall, we show that lncRNAs are dynamically regulated upon EBOV infection suggesting that they play important roles in the host response to EBOV infection in a time and cell-type specific manner.

### LncRNAs are involved in the host innate immune response to EBOV

Although we detected many lncRNAs that change expression upon EBOV infection, the large majority of those remain functionally uncharacterized. Some lncRNAs exert their modulatory role in a *cis*-regulatory fashion (Gil and Ulitsky 2019). To identify possible *cis*-regulatory lncRNAs, we first detected 76 lncRNAs protein-coding genes pairs that were both differentially expressed in the same cell type and in close physical proximity (<1Mbp). From those, however, only 17 pairs were significantly correlated (Spearman correlation test, P-value < 0.05, Supplemental Fig. S7). Thus, although we found pairs of lncRNA and protein-coding genes co-located and co-expressed at cell-type resolution, we did not observe them to be significantly co-located more often than expected by chance (Fisher’s exact test, OR = 0.44, P-value > 0.05) (see Methods).

To explore further the pathways and putative functions of our DE lncRNAs, we built a cell-type specific co-expression network in monocytes (see Methods) and identified seven modules. The three largest modules were enriched in different functions related to the innate immune response (Figure 4A, Supplemental Fig. S8A-E). Module 1 was enriched in “regulation of response to stress” and was mostly composed of downregulated genes (Figure 4A, Supplemental Fig. S8C, Supplemental Table S3). Cellular stress and subsequent inhibition of translation is a common host defence response to viral infections (McCormick and Khaperskyy 2017; Walsh et al. 2013) and might reach its peak at the late stage of the infection when the amount of infected cells is highest. Consistent with this, most of the genes in the module showed the strongest expression changes at late stages (Supplemental Fig. S8A). Also, one of the lncRNAs in this module, MIR22HG, has been reported to be sensitive to expression changes upon cellular stress (Tani and Torimura 2013). The second largest module was related to “innate immune response” (Figure 4A, Figure S8D, Supplemental Table S3). Importantly, it includes many interferon-stimulated genes (ISG) such as MX1, IFIT2 and ISG15. These ISGs are all up-regulated at early and middle stages of infection (as expected)(Kotliar et al. 2020)(Figure 4A, Supplemental Fig. S8A-B), as is the lncRNA ENSMMUG00000064224 directly connected to them in the network. Similarly to ISG, ENSMMUG00000064224 changes its expression in B and T cells as well as in monocytes (Figure 3G, Supplemental Fig. S8B), supporting the hypothesis that it may play an ISG-like role upon infection that is shared across cell types (Kotliar et al. 2020). Module 3, had most genes downregulated and was enriched in purine metabolism (Figure 4A, Figure S8E, Supplemental Table S3). This is consistent with the reported decreased nucleotide availability interfering with viral replication as an anti-viral host response to EBOV infection (Luthra et al. 2018).

The remaining modules did not have significant functional enrichments likely due to their smaller sizes. However, all except one had central regulators or downstream effectors of the innate immune response (Supplemental Table S3). For example, the smallest module harboured ETS1, a transcription factor that controls the expression of cytokine and chemokine genes, together with one of its main targets, ILR7 (Grenningloh et al. 2011).

Overall, our co-expression network analysis has allowed assigning some of our DE lncRNAs to specific functional roles related to innate immune processes such as interferon response, stress response, and purine metabolism.

### LncRNAs are up and down-regulated upon viral entry and replication in infected monocytes

Upon entry into the cell and replication, EBOV hijacks some of the cells’ defenses by down-regulating anti-viral genes and up-regulating pro-viral genes (Kotliar et al. 2020). To study whether lncRNA expression is also subject to changes upon viral replication and may be therefore part of pathways that EBOV leverages to attack the host cell, we tested for an association between viral load and lncRNA expression in infected monocytes. We used macaque PBMCs infected *ex vivo* (Figure 5A, Supplemental Fig. S9A-E), as studying the *ex vivo* experimental setup allowed for higher viral exposure and consequently a higher number of infected cells. We identified 16 lncRNAs significantly correlated with viral load (Spearman correlation test, q-value < 0.05) (Supplemental Table S4), the majority of which (12) were positively correlated (Figure 5B).

**Figure 5.**
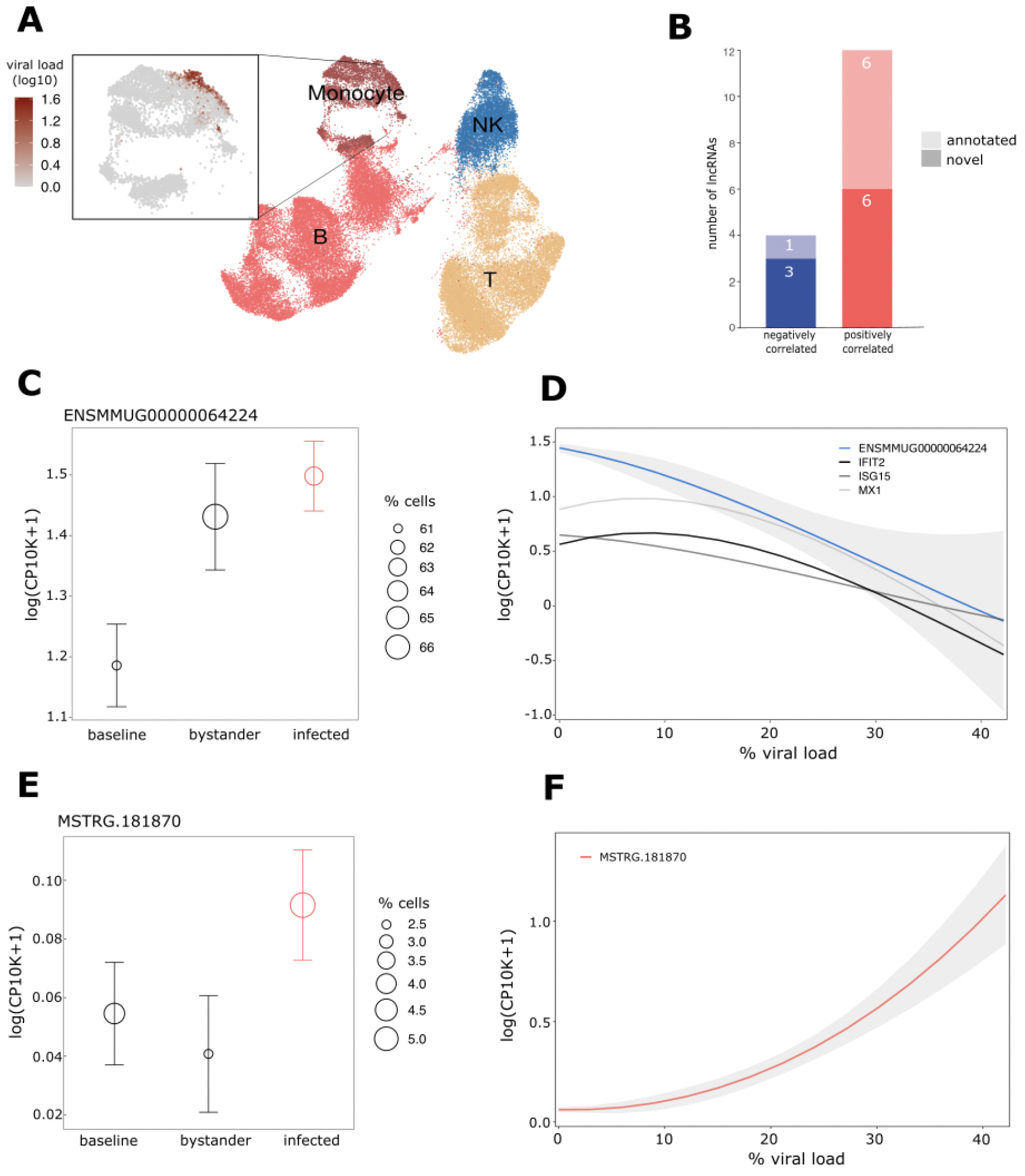
LncRNAs undergo expression changes exclusively in infected monocytes. **(A)** UMAP embedding of 56,317 cells from the *ex vivo* dataset. Cell types are indicated by different colours. The magnified UMAP on the left highlights the cells with viral transcripts. **(B)** Number of novel and annotated lncRNA positively and negatively correlated with viral load. **(C)** Expression of ENSMMUG00000064224 in baseline as well as bystander and infected cells at 24h post-infection. Marker: mean; error bars: 95% confidence interval. Dot sizes represent the percentage of cells in which the gene is expressed in each cell group. **(D)** ENSMMUG00000064224, IFIT2, ISG15 and MX1 expression changes with viral load. **(E)** Expression of MSTRG.181870 in baseline, bystander and infected cells at 24h post-infection. Marker: mean; error bars: 95% confidence interval. Dot sizes represent the percentage of cells in which the gene is expressed in each cell group. **(F)** MSTRG.181870 expression changes with viral load.

ENSMMUG00000058644 and MSTRG.15458, which had the strongest correlations, were also significantly correlated at nominal P-values in the *in vivo* dataset (Spearman ρ= 0.10, P-value=0.03 and Spearman ρ= −0.12, P-value=0.01 respectively), suggesting that the *in vivo* dataset might not have enough infected cells to identify significant correlations. In line with this, lncRNAs correlated with viral load were expressed in significantly fewer cells *in vivo* compared to *ex vivo* (Mann-Whitney U test, P-value < 2 x 10^−10^).

Eleven out of the 16 identified lncRNAs had not been detected as differentially expressed in monocytes (Figure 3C, Supplemental Table S1, S4) suggesting that most of these lncRNAs change expression exclusively in infected cells, and it highlights the power of the single-cell analysis. The remaining 5 lncRNAs had been identified as differentially expressed in monocytes upon infection. Strikingly though, lncRNAs upregulated during infection in the general monocyte population were negatively correlated with the viral load in infected cells. Conversely, lncRNAs downregulated during infection in monocytes increased their expression with viral load in infected monocytes (Figure 6C-F, Supplemental Fig. 10A-F). This suggests that such lncRNAs are part of changes directly triggered by viral cellular entry and replication.

The lncRNA with the strongest drop in expression upon increased viral load was ENSMMUG00000064224 (Spearman ρ= −0.11, q =6.50 · 10^−5^) (Figure 5C, Supplemental Table S1, S4). We previously classified ENSMMUG00000064224 as an interferon response lncRNA based on our co-expression network analysis (Figure 4A). In accordance with this, ENSMMUG00000064224 is upregulated upon infection but is negatively correlated with viral load similarly to known ISGs such as IFI2, ISG15 and MX1 (Figure 5D). (Kotliar et al. 2020). Previous work suggested that EBOV actively downregulates the interferon response pathway to allow viral replication. Our results suggest that some lncRNAs may also be hijacked by EBOV to promote viral entry and replication.

## Discussion

Despite the growing evidence associating lncRNAs to crucial immune regulatory roles (Heward and Lindsay 2014; Marina R. Hadjicharalambous 2019), studies focusing on the long non-coding RNA response upon infection at single-cell resolution are scarce (Li et al. 2020a; Wang et al. 2020). A main reason for this is that lncRNA annotations remain largely incomplete. Here, we annotated ~4,000 novel lncRNAs, which almost doubled the size of the current long non-coding annotation and described essential properties that distinguish lncRNAs from protein-coding genes. ~500 novel lncRNAs are conserved in human; as they may play biologically comparable functions in both species, our findings may have direct implications for EBOV infection in humans. In addition, we identified lncRNAs that are up and down-regulated upon infection in a cell-type and disease-stage specific manner and we were able to pinpoint the pathways they are related to, including interferon-mediated immune response and host metabolism. Overall, our expanded lncRNA annotation together with our in-depth characterization will serve as a reference atlas of macaque lncRNAs for future studies.

Our results suggest that a main differential property of lncRNAs compared to protein-coding genes is their expression in fewer cells. More importantly, when controlling for the number of cells in which lncRNAs are expressed, lncRNAs have similar expression levels, cell-type specificity, and dynamic regulation compared to protein-coding genes. Liu et al. made a similar observation in brain tissue although their study was heavily constrained by the number of cells (<250 cells). This result raises the following question: why do lncRNAs have expression that is restricted to few cells? Previous work showed that lncRNAs have fewer transcription factor binding sites and higher chromatin repressive marks in their promoter regions compared to equally expressed protein-coding genes (Melé et al. 2017). In addition, transcription factor binding sites in lncRNA promoters are less complex than those in protein-coding genes suggesting that fewer transcription factors can bind to a specific binding site in a lncRNA promoter than in a protein-coding promoter (Mattioli et al. 2019). Our results are consistent with a model in which promoters of lncRNAs differ from those of equally expressed protein-coding genes in the probability of engaging in active transcription rather than in the strength of the transcriptional response.

Additionally, the novel metric Upsilon which we designed to study the cell-type specificity may have wide use in future scRNA-Seq studies. Compared to the alternative metric Tau, repurposed from bulk analyses, Upsilon is superior in discriminating markers from housekeeping genes as well as genes with intermediate cell-type specificity patterns.

Our single-cell transcriptomic analysis allowed us to identify cell type specific lncRNA expression changes correlated with viral load. Recent work has shown that EBOV may induce host gene expression changes to its benefit, by up- and down-regulating pro- and anti-viral genes respectively (Kotliar et al. 2020). Our identified lncRNAs could be part of pathways hijacked by EBOV to efficiently attack the host cell, suggesting that these genes play crucial roles during the host immune response. However, a major limitation of our analysis is the number of cells we could analyze as the number of infected cells where we found lncRNAs expressed is limited, especially *in vivo* as very few cells are infected even in severely diseased animals. We speculate that further studies with a higher amount of infected cells will allow for a more comprehensive detection of lncRNAs changing their expression upon EBOV infection.

In summary, this study sheds light on the roles of lncRNAs in response to EBOV infection and paves the way for future studies on how to systematically analyze lncRNAs at single-cell resolution.

## Material and Methods

### RNA sample processing

For *de novo* annotation, we performed paired-end, strand-specific bulk RNA-sequencing (RNA-Seq) on high-quality, commercially available rhesus monkey total RNA (Zyagen, San Diego CA, USA; hereafter referred to as Zyagen) of non-infected samples from 10 different tissues (Supplemental Table S1). Briefly, we depleted ribosomal RNA and performed random-primed cDNA synthesis (Matranga et al. 2014), followed by second strand marking and DNA ligation (Levin et al. 2010) with adaptors containing unique molecular identifiers (UMIs) (MacConaill et al. 2018)(IDT, Coralville IA, USA). We performed a similar bulk RNA-Seq protocol without UMIs on rhesus monkey RNA samples from the study by (Luke et al. 2018). Additionally, we downloaded bulk RNA-Seq from the NCBI Gene Expression Omnibus (GEO; http://www.ncbi.nlm.nih.gov/geo/) from the accession number GSE115785. For the single-cell RNA-Seq analysis, we downloaded the dataset from the NCBI Gene Expression Omnibus (GEO; http://www.ncbi.nlm.nih.gov/geo/) with accession number GSE158390.

### QC and mapping

First, we merged Ensembl Mmul_10 release 100 assembly and Ensembl release 100 gene annotation with the Ebola virus/H. sapiens-tc/COD/1995/Kikwit-9510621 (GenBank #KU182905.1; *Filoviridae: Zaire ebolavirus*) assembly and annotation respectively, and used them throughout all downstream analyses. We used Hisat v2.1.0 (Kim et al. 2019) to compute assembly indexes and known splice sites and mapped each samples’ reads to the merged assembly. We ran Hisat2 with default parameters, except for rna-strandness, which we set according to the experiments’ strandness (Supplemental Table S1), previously inferred with InferExperiment.py from RSeQCc v3.0.0 (Wang et al. 2012). We sorted mapped bam files with samtools sort v1.9 (Li et al. 2009) with default parameters. We retained only paired and uniquely mapped reads using samtools view with parameters *-f3 -q 60*. Additionally, we removed duplicates from the samples tagged with UMIs (Zyagen) (Supplemental Table S1) with umi_tools dedup v1.0.0 (Smith et al. 2017). We excluded all samples with less than 10M sequenced reads, a mapping rate lower than 0.3 or a genic mapping rate lower than 0.7. We defined the genic mapping rate as the proportion of exonic and intronic reads, as computed by read_distribution.py from RSeQCc v3.0.0 (Wang et al. 2012) (See Supplemental Table S1).

### LncRNA discovery pipeline

We ran *de novo* transcriptome assembly separately on each sample with Stringtie v1.3.6 (Pertea et al. 2015), with default parameters except for strand information that was set depending on the dataset (Supplementary Table 1). We used Stringtie to merge all the *de novo* assemblies using the parameter “--*merge*”. To identify novel transcripts absent from the reference annotation, we used Gffcompare v0.10.6 and retained exclusively the transcripts with class code “u” and “x”, corresponding to intergenic and antisense transcripts. We removed mono-exonic transcripts, transcripts shorter than 200 bp, and transcripts not expressed (log(TPM) < 1) in at least three samples. To assess the coding potential of the newly assembled transcripts, we used three sequence-based lncRNAs prediction tools: Coding Potential Assessment Tool (CPAT)(Wang et al. 2013), Coding Potential Calculator (CPC2)[(Kang et al. 2017)] and Coding-Non-Coding Identifying Tool (CNIT)(Guo et al. 2019) with default parameters. For each prediction tool independently, we removed genes with at least one isoform predicted as non-coding and one predicted as protein-coding. We considered a gene to be a long non-coding RNA if the three tools classified it as non-coding. We then merged the obtained list of novel lncRNAs to the reference annotation and used it in downstream analyses. To benchmark our lncRNAs discovery pipeline, we predicted the biotype of annotated genes (Ensembl v100) (coding or non-coding) and compared our predictions to their annotated biotype. To compare lncRNA and protein-coding transcript length, number of exons and exon length, we considered the longest transcript per gene. To identify lncRNAs orthologs to human, we used the synteny-based lncRNAs detection tool slnky v1.0 on human hg38 assembly and gencode hg38 v23 annotation (Chen et al. 2016). For the sake of reproducibility, the lncRNAs discovery pipeline is implemented in Nextflow (Di Tommaso et al. 2017) and combined with Singularity software containers.

### Single-cell RNA sequencing data and processing

We used two publicly available single-cell RNA-Seq datasets of macaque PBMCs infected with EBOV i*n vivo* and *ex vivo* (Kotliar et al. 2020). For the *in vivo* dataset, samples were extracted at different days post-infection (38,067 cells). In the *ex vivo* dataset, PBMCs were divided into three batches after sampling. Live virus was added (MOI of 0.1 pfu/cell) to the first batch, hereafter referred to as live; inactivated virus, which was made unable to replicate through gamma irradiation, was added to the second one, hereafter referred to as irrad; no addition was made to the third batch which was established as control, hereafter referred to as media. Each batch was sampled twice, at 4 and 24 hours, and submitted to scRNA-Seq, performed with Seq-Well (48,350 cells). To obtain gene expression quantification, we implemented the Drop-seq analysis pipeline (https://github.com/broadinstitute/Drop-seq) in the form of a Nextflow (Di Tommaso et al. 2017) pipeline combined with Singularity containers for the sake of reproducibility. We followed the best practices for single-cell analyses as described by (Luecken and Theis 2019). to manually identify suitable filtering thresholds. For the in vivo dataset, we selected cells with at least 1,000 and a maximum of 10,000 UMIs, at least 600 and a maximum of 2,000 detected genes. Additionally, we excluded cells with more than 5% of mitochondrial reads. In the *ex vivo* dataset, PBMCs were infected with a higher EBOV dose than in vivo, resulting in an increased number of infected cells and also, an increased number of viral transcripts within infected cells. To make sure we did not exclude highly infected cells from our analysis, we increased the upper thresholds for the number of allowed UMIs per cell to 15,000 and the number of genes detected per cell to 4,000.

Additionally, in both datasets, we excluded genes expressed in fewer than 10 cells. We used Scrublet v.0.2.1 (Wolock et al. 2019) for doublet detection. In the *in vivo* dataset, we applied the IntegrateData method of Seurat v3.0 (Stuart et al. 2019) that uses canonical correlation analysis (CCA) to correct for fresh versus frozen batch effect. We normalized counts to log(CP10K+1), the default in the Seurat package. To replicate some of our observations in a human dataset, we used available gene counts of human healthy PBMCs from 10xGenomics (https://www.10xgenomics.com/resources/datasets/33-k-pbm-cs-from-a-healthy-donor-1-standard-1-1-0) (32,738 available cells) and human Ensembl version 100 gene annotation.

### Single-cell clustering and cell-type identification

To cluster cells, we used the Louvain algorithm as implemented in the Seurat package (Stuart et al. 2019). To identify cluster-specific genes, we ran a differential expression analysis between each cluster and all the remaining ones using the Seurat function FindAllMarkers. Based on the expression levels of known marker genes, we classified clusters into specific cell types using (Supplemental Fig. 3A, Supplemental Fig. S3A).

### LncRNA and protein coding gene comparisons

We used Seurat’s normalization values (log(CP10K+1)) to compare expression levels between lncRNAs and protein-coding genes. We considered a gene to be expressed in a cell when its normalized expression value was larger than 1. Median expression values were calculated exclusively across cells in which the gene was expressed. We used the MatchIt R package (https://www.rdocumentation.org/packages/MatchIt/) to obtain the pairs of lncRNA and protein-coding genes matched either by median expression or by the percentage of cells in which they were expressed.

### Cell-type specificity estimates

We considered two distinct cell-type specificity measurements. First, we leveraged Tau (Kryuchkova-Mostacci and Robinson-Rechavi 2017), a metric originally designed to assess tissue-specificity. Instead of calculating for each gene the mean expression per tissue, we calculated mean expression per cell type, including zeros. Tau was calculated as follows:

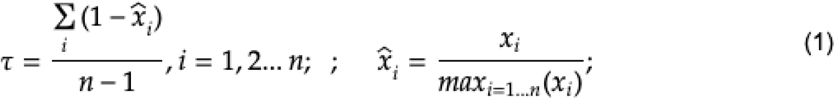

where *x_i_* is the mean expression of a gene in cell-type *i* and *n* is the total number of cell types. Additionally, we designed a score (Upsilon, ν) that relies purely on the proportion of cells in which each gene is expressed, which was calculated as follows:

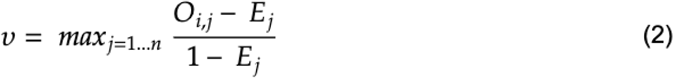

where:

*O_ij_* is the observed proportion of cells in which gene *i* is found expressed in cell-type *j*. To calculate the proportions of cells in which each gene is expressed per cell type, we considered only the cells in which we detected the gene as expressed, so that, per gene, the proportions assigned to the different cell-types sum up to one.
*E_ij_* is the expected proportion of cells in which gene *i* would be expressed in cell-type *j* if it was not cell-type specific. The expected proportion of cells for cell type *j* is equal for all the genes and corresponds to the proportion of cells of cell-type *j* in the dataset.

Then, we divided the difference between the observed and expected proportions by the maximum value this difference could reach. The maximum value is reached when the gene is expressed in all cells of one cell type, which is the difference between 1 and the expected proportion. The value, therefore, ranges from 0 to 1. We then calculated the specificity of each gene to each of the cell types and reported the maximum of these values as the gene’s global specificity score.

To explore the behaviour of the two cell-type specificity metrics, we simulated different scenarios for genes with three different levels of cell-type specificity (highly, intermediate or lowly cell-type specific genes), in a dataset with three different cell types in different proportions (50%, 30% and 20% of the total number of cells) (Supplemental Fig. S4).

### Differential expression analysis

We grouped samples of the *in vivo* dataset into baseline, early, middle and late stages, based on the day post-infection of the sample (Figure 2A). We performed differential expression analysis comparing each stage of the infection(early, middle, late) to baseline in Monocytes, B and T cells separately. We excluded neutrophils because they were detected in PBMCs exclusively at later stages of infection. We adopted the statistical framework MAST (Finak et al. 2015), using normalized UMI counts as input. As covariates, we used the number of genes detected per cell and a variable corresponding to whether the sample had been frozen or not. We used Fisher’s exact test to investigate whether lncRNAs have differential expression patterns more cell-type specific than protein-coding genes. The two tested variables are gene biotype and whether the gene is DE in one or more cell types.

### Gene colocation analysis

We used the GenomicRanges package (https://bioconductor.org/packages/release/bioc/html/GenomicRanges.html) to calculate the genomic distance between genes in the macaque Ensembl v100 annotation. To test whether DE lncRNAs were closer to DE protein-coding genes more often than not DE lncRNAs, we set up a Fishers’ exact test. The two tested variables were whether the lncRNA is DE and whether is it in cis to a DE protein-coding gene.

### Co-expression network

We built a co-expression network using all differentially expressed genes in monocytes with GrnBoost2 (Moerman et al. 2019). To focus on the co-regulatory network involving lncRNAs, we only retained edges connected to at least one lncRNA. Also, we retained exclusively the top 0.5% edges, when sorted by weight. We identified communities with the Louvain algorithm (Blondel et al. 2008) and reported those with at least 5 edges. For the functional enrichment of the modules, we used the R package clusterProfiler v4.2.0(Wu et al. 2021).

### Correlation with viral load

To identify genes whose expression correlates with viral transcripts changes in the cell, we only considered infected monocytes at a late stage of infection (24h post-infection *ex vivo*, days post-infection 5-8 *in vivo*). We computed the spearman correlation coefficient between the viral load (log10) and each gene’s normalized expression (log(CP10K+1)). Normalized expression values were calculated after removing viral transcripts to avoid library size normalization biases. P-values were corrected with Benjamin and Hochberg multiple testing correction (Benjamini and Hochberg 1995).

### Data access

Sequencing data has been submitted to the NCBI Gene Expression Omnibus (GEO; http://www.ncbi.nlm.nih.gov/geo/) under accession number GSE192447. All pipelines and scripts used in this study are available at https://gitlab.bsc.es/melebsc/ebola.

## Competing interest statement

The authors declare no competing interests.

## Acknowledgements

We thank Kaia Mattioli for thoughtful comments on the manuscript and Aida Ripoll-Cladellas for feedback and helpful discussions. This material was based upon work supported by the Ramon y Cajal RYC-2017-22249 grant, the Howard Hughes Medical Institute Investigator Award (P.C.S), the National Institute of Allergy and Infectious Diseases (NIAID) U19AI110818, the US Food and Drug Administration (FDA) contract HHSF223201810172C. We acknowledge SAB Biotherapeutics as partners for providing study materials from the study by (Luke et al. 2018) and for their collaborative support that allowed the study success.

## Author contributions

L.S. performed most computational analysis. M.S.R. performed viral load’s correlation computational analysis. M.M. designed the project. J.L.R contributed to the study design. M.M and R.G.P supervised the analysis. L.S., M.M. and R.G.P wrote the manuscript. A.E.L., G.C.A, K.J.S. and S.W. did all experimental work. F.R. contributed to the design of the cell-type specificity score. L.E.H., R.S.B and P.C.S. designed and led all experimental work. All authors have read and approved the manuscript for publication.

## Notes

### Competing Interest Statement

The authors have declared no competing interest.

