## Supplemental Materials for "*In vivo* single-cell profiling of lncRNAs during Ebola virus infection"

**Table of Contents:**

**Supplemental Figure S1.** Identification and characterization of novel lncRNAs.

**Supplemental Figure S2.** Novel intergenic and novel antisense lncRNAs resemble annotated lncRNAs.

**Supplemental Figure S3.** Levels and proportion of cells of lncRNA and protein-coding genes expression at single-cell resolution.

**Supplemental Figure S4:** Cell-type specificity scores behaviour across simulated scenarios.

**Supplemental Figure S5.** Cell-type specificity scores and proportion of cells in which lncRNAs and protein-coding genes are expressed.

**Supplemental Figure S6.** Differential expression patterns of lncRNAs upon EBOV infection.

**Supplemental Figure S7**. Characterization of *cis*-regulatory activity of lncRNAs.

**Supplemental Figure S8.** Co-expression network of lncRNAs and protein-coding genes upon EBOV infection in monocytes.

**Supplemental Figure S9.** *Ex vivo* dataset processing.

**Supplemental Figure S10.** LncRNAs undergo expression changes within infected monocytes.

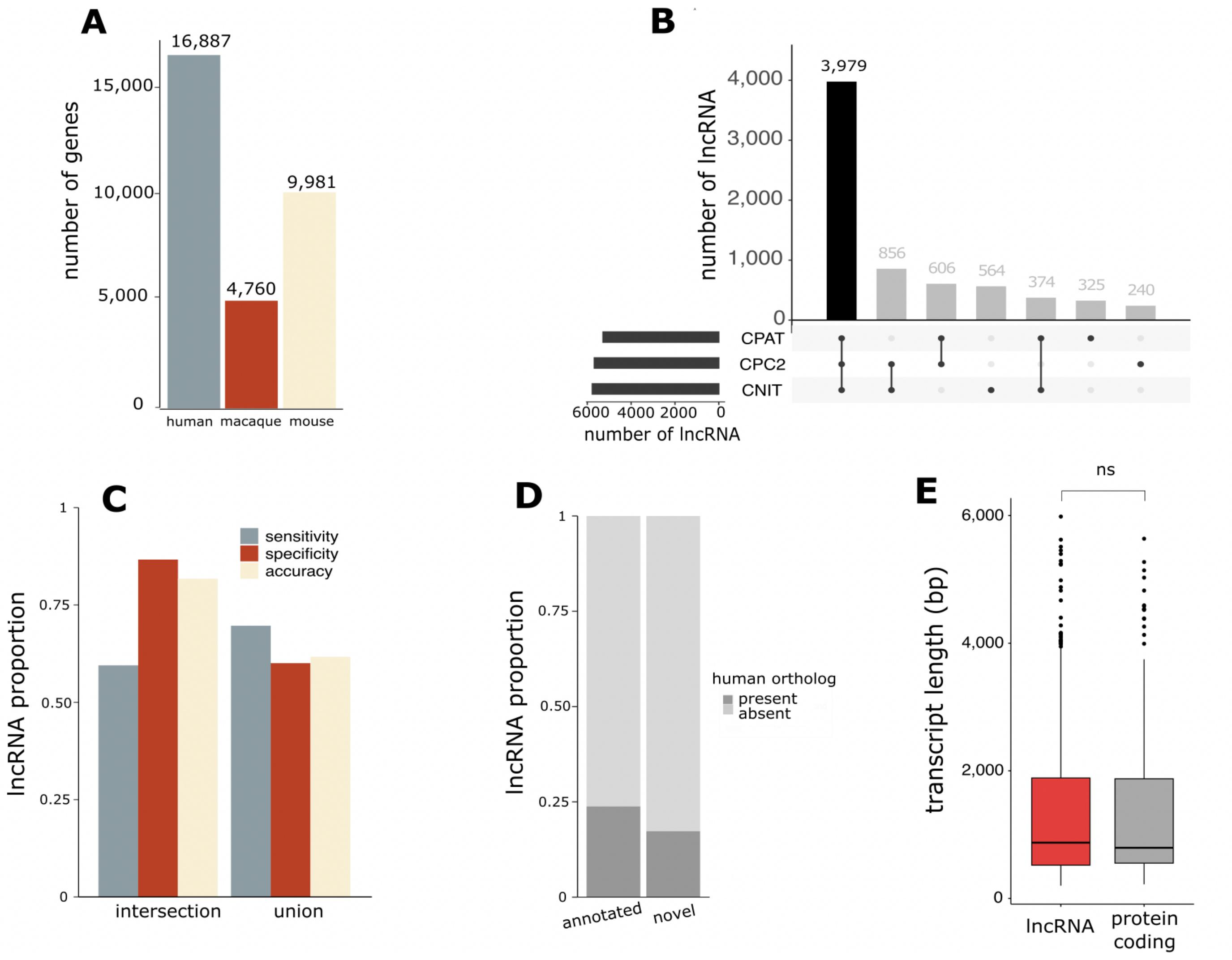


**Supplemental Figure S1. Identification and characterization of novel lncRNAs. (A)** Number of annotated lncRNA genes in human, macaque and mouse in the Ensembl release 100. **(B)** Overlap of the genes predicted as non-coding by the CPAT, CPC2 and CNIT tools. Highlighted in black is the intersection of the prediction of the three tools, which represents the set we used for downstream analyses. **(C)** Benchmarking measures for non-coding genes obtained by the intersection (left) and union (right) of biotype predictions from CPAT, CPC2 and CNIT. **(D)** Proportion of genes for which we identified a human ortholog in annotated (left) and novel lncRNAs (right). **(E)** Transcript length of the novel lncRNAs and their corresponding human ortholog (Paired Wilcoxon signed-rank test, P-value >0.05).

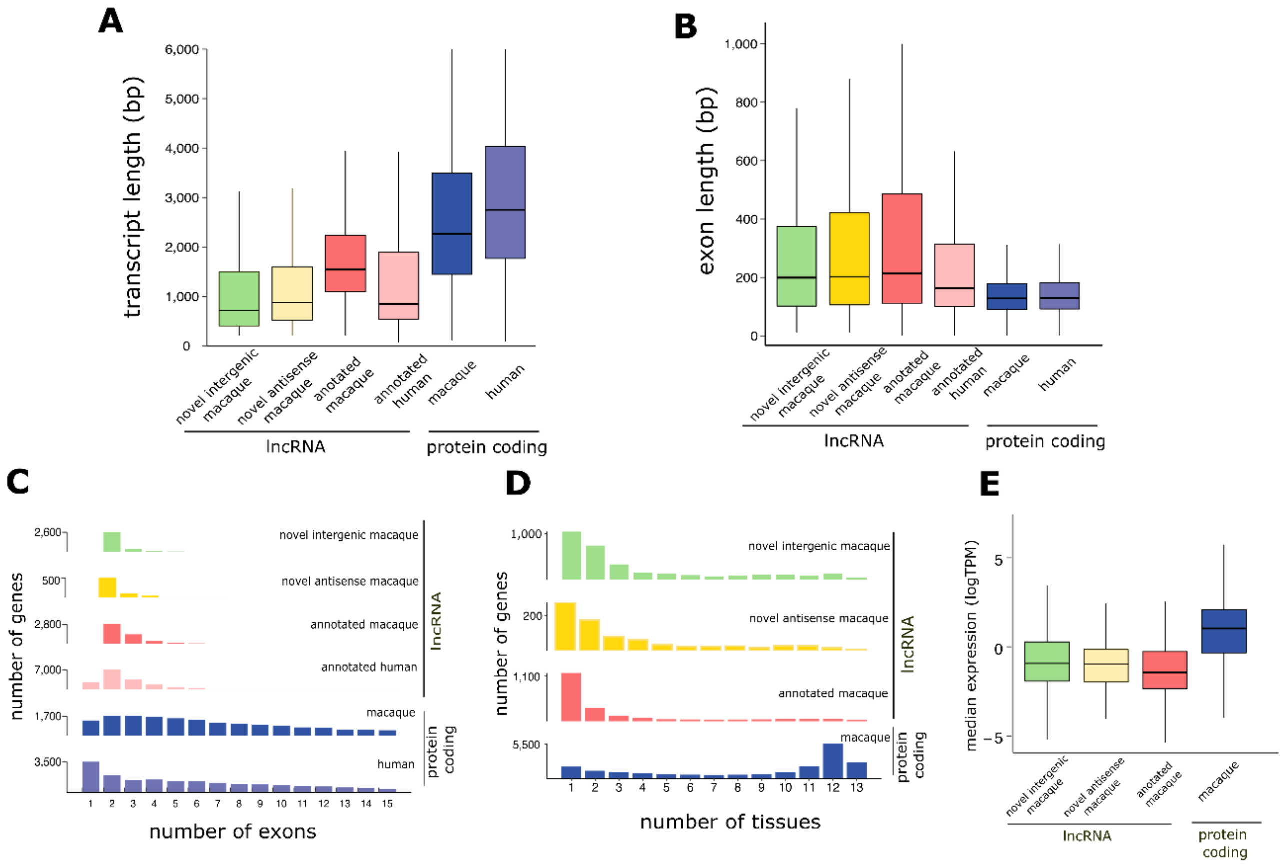


**Supplemental Figure S2. Novel intergenic and novel antisense lncRNAs resemble annotated lncRNAs.** (**A**) Distribution of transcripts length (**B**), exons length (**C**), number of exons per transcript (**D**), number of tissue in which genes show expression (**E**) and median expression levels (**F**) of novel intergenic lncRNA (green), novel antisense lncRNA (red), and previously annotated macaque (red) and human (blue) lncRNA.

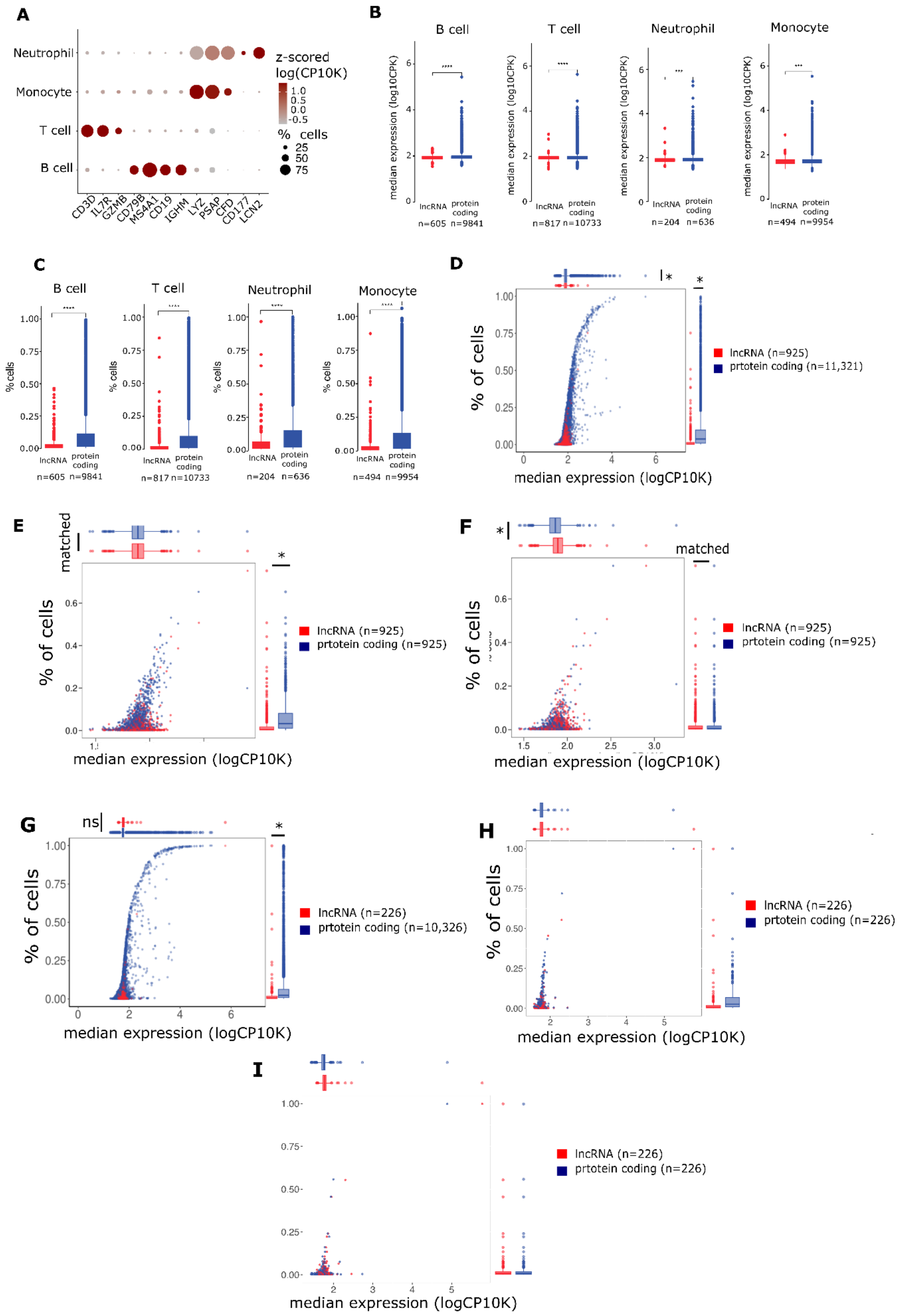


**Supplemental Figure S3. Levels and proportion of cells of lncRNA and protein-coding genes**

**expression at single-cell resolution. (A)** Dot plot showing the expression of cell-type markers which were used for cell type assignment in the *in vivo* dataset. Dot's colours represent the expression level (z-scored log(CP10K)) of the gene in each cell type. Dot's sizes represent the percentage of cells in which the gene is detected as expressed per cell type. **(B)** Median expression levels and **(C)** percentage of cells in which each gene is expressed, across cell-types and separated by gene classes. Asterisk indicates level of significance (Mann Whitney U test: ** P-value < 0.01, *** P-value < 0.001). **(D)** Scatter plot displaying the median expression level of lncRNAs (red) and protein-coding genes (blue) versus the proportion of cells in which they are expressed **(E)** Same as D but for lncRNAs and protein-coding genes matched by median expression levels. **(F)** Same as D but for lncRNAs and protein-coding genes matched by the percentage of cells in which they are expressed. **(G)** same as D but for lncRNAs and protein-coding genes in human PBMCs. **(H)** Same as D but for lncRNAs and protein-coding genes in human PBMCs matched by median expression levels. **(I)** Same as D but for lncRNAs and protein-coding genes in human PBMCs matched by the percentage of cells in which they are expressed.

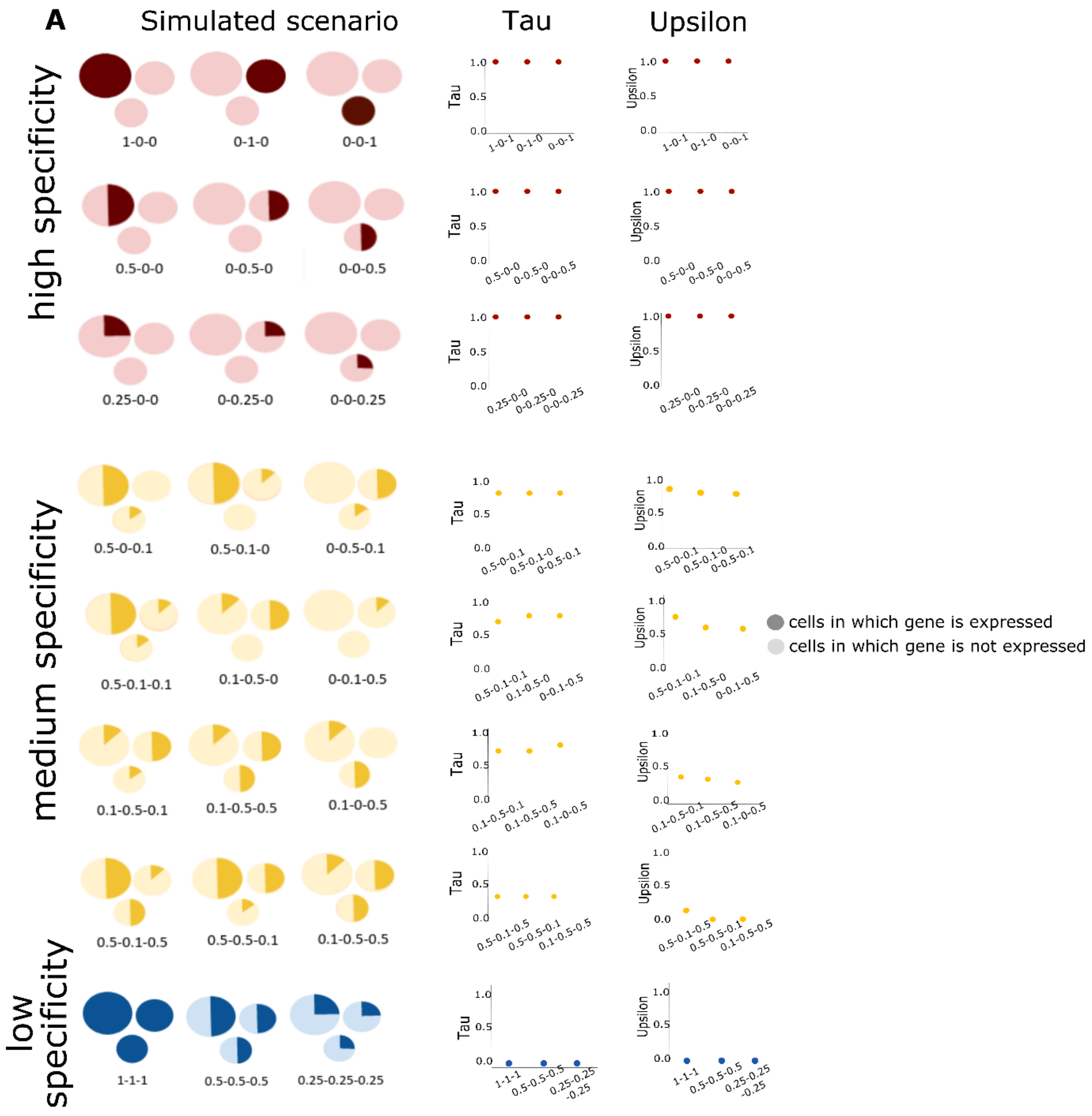


**Supplemental Figure S4. Cell-type specificity scores behaviour across simulated scenarios.**

**(A)** Schematic representation of the simulated scenarios where we calculate the Tau and Upsilon scores for genes with high, intermediate or low cell-type specific (red, yellow and blue respectively). Cells are split into three cell types, which represent 50%, 30% and 20% of the total number of cells. Proportions highlighted in darker and lighter colours represent cells in which the gene is expressed and not expressed respectively. Numbers below each plot correspond to the proportion of cells in which the gene is expressed, sorted from the cell types with the highest proportion of cells to the lowest proportion of cells. Highly specific genes can be expressed in 100%, 50% or 25% of the cells of exclusively one cell type. Genes with intermediate cell-type specificity can be (1) expressed in 50% of the cells of one cell-type, 10% of a second cell-type and show no expression in the remaining cell-type, (2) expressed in 50% of the cells of one cell-type and 10% in all other cell types, or (3) expressed in 50% of the cells of two cell types and not expressed in the remaining cell-type. Lastly, low cell-type specific genes can be expressed in 100%, 50% and 10% of all cell types. Corresponding **(B)** Tau and **(C)** Upsilon cell-type specificity scores.

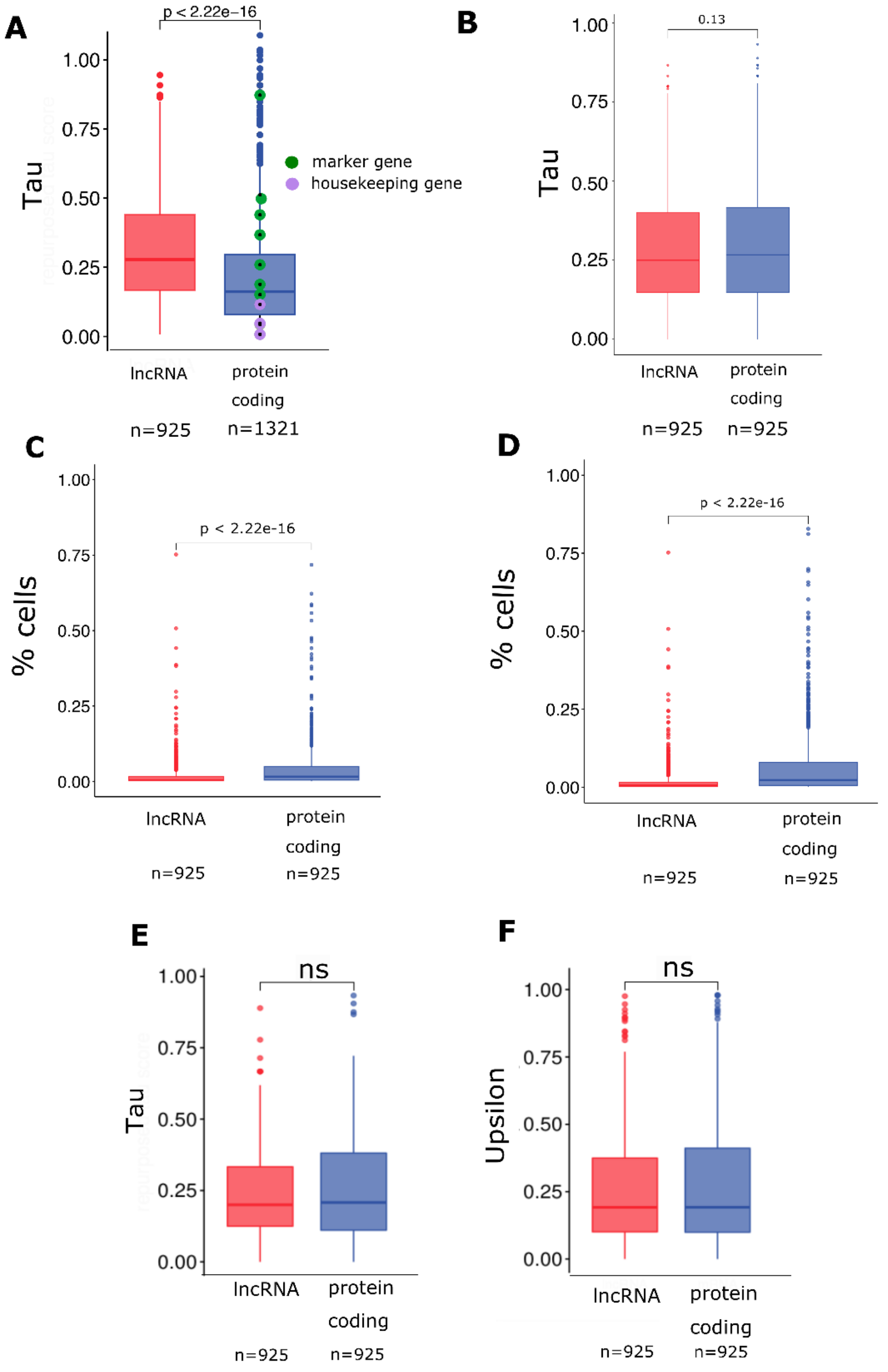


**Supplemental Figure S5. Cell-type specificity scores and proportion of cells in which lncRNAs and protein-coding genes are expressed. (A)** Tau specificity scores calculated on the *in vivo* dataset, separated between lncRNAs (red) and protein-coding genes (blue). Cell-type marker genes are highlighted in green, housekeeping genes in purple. **(B)** Cell-type specificity scores (Tau) of lncRNAs and protein-coding genes matched by the percentage of cells in which they express. Same as B when lncRNAs and protein-coding genes are matched by Tau **(C)** and Upsilon **(D)** cell-type specificity scores. Tau **(E)** and Upsilon specificity score **(F)** calculated for the human healthy PBMCs dataset.

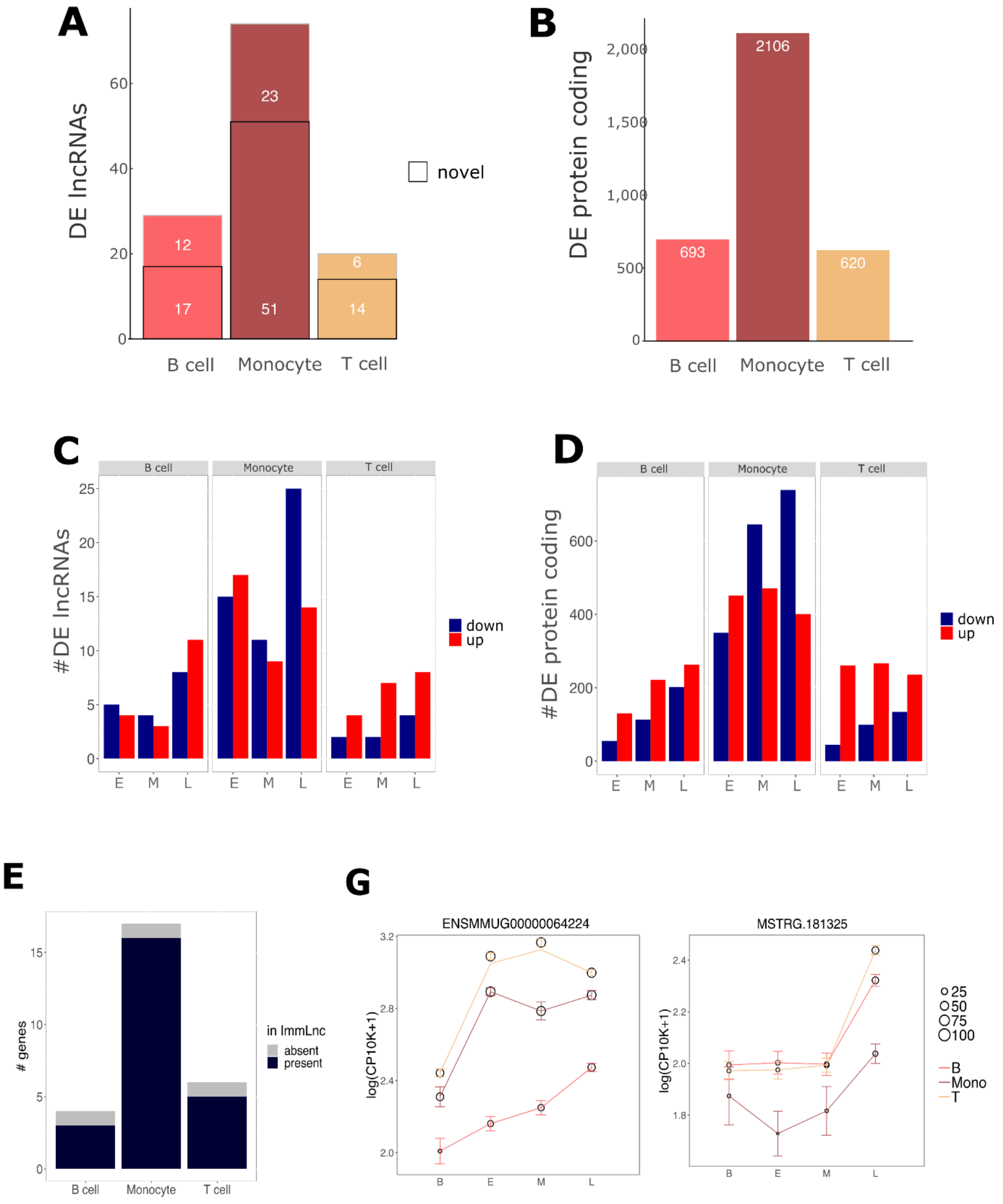


**Supplemental Figure S6. Differential expression patterns of lncRNAs upon EBOV infection.**

**(A)** Bar plot showing the number of differentially expressed lncRNAs detected upon EBOV infection *in vivo*, coloured by cell type and separated by whether they were annotated (no border) or identified through *de novo* annotation (black border). **(B)** Bar plot showing the amount of differentially expressed protein-coding genes detected upon EBOV infection *in vivo* coloured by cell-type. Bar plot showing the number of differentially expressed lncRNAs **(C)** and protein coding genes **(D)** detected upon EBOV infection *in vivo*, separated by the directionality of the dysregulation. **(E)** Bar plot showing the number of differentially expressed lncRNAs detected upon EBOV infection *in vivo* for which we could identify an ortholog. Colour code indicates whether the lncRNA had been previously reported in ImmLnc. **(G)** Expression patterns of the lncRNAs ENSMMUG00000064224 and MSTRG.181325 at different stages of infection in monocytes, B and T cells. Marker: mean; error bars: 95% confidence interval. Dot's sizes represent the percentage of cells in which the gene is expressed in each cell type.

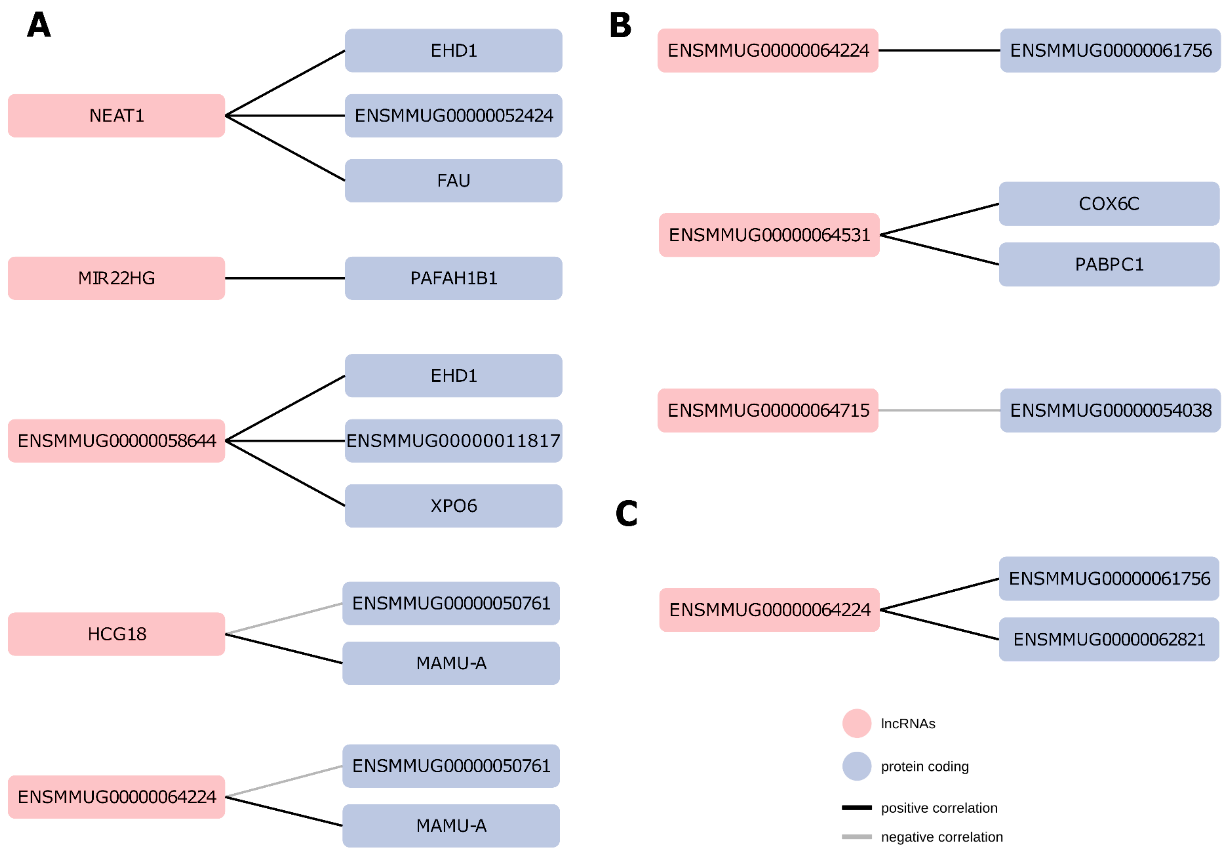


**Supplemental Figure S7. Characterization of *cis*-regulatory activity of lncRNAs.** Neighbouring lncRNAs (red) and protein-coding genes (blue) differentially expressed and significantly correlated (Spearman correlation test, P-value < 0.05) upon EBOV infection *in vivo* in Monocytes (**A**), T (**B**) and B (**C**) cells.

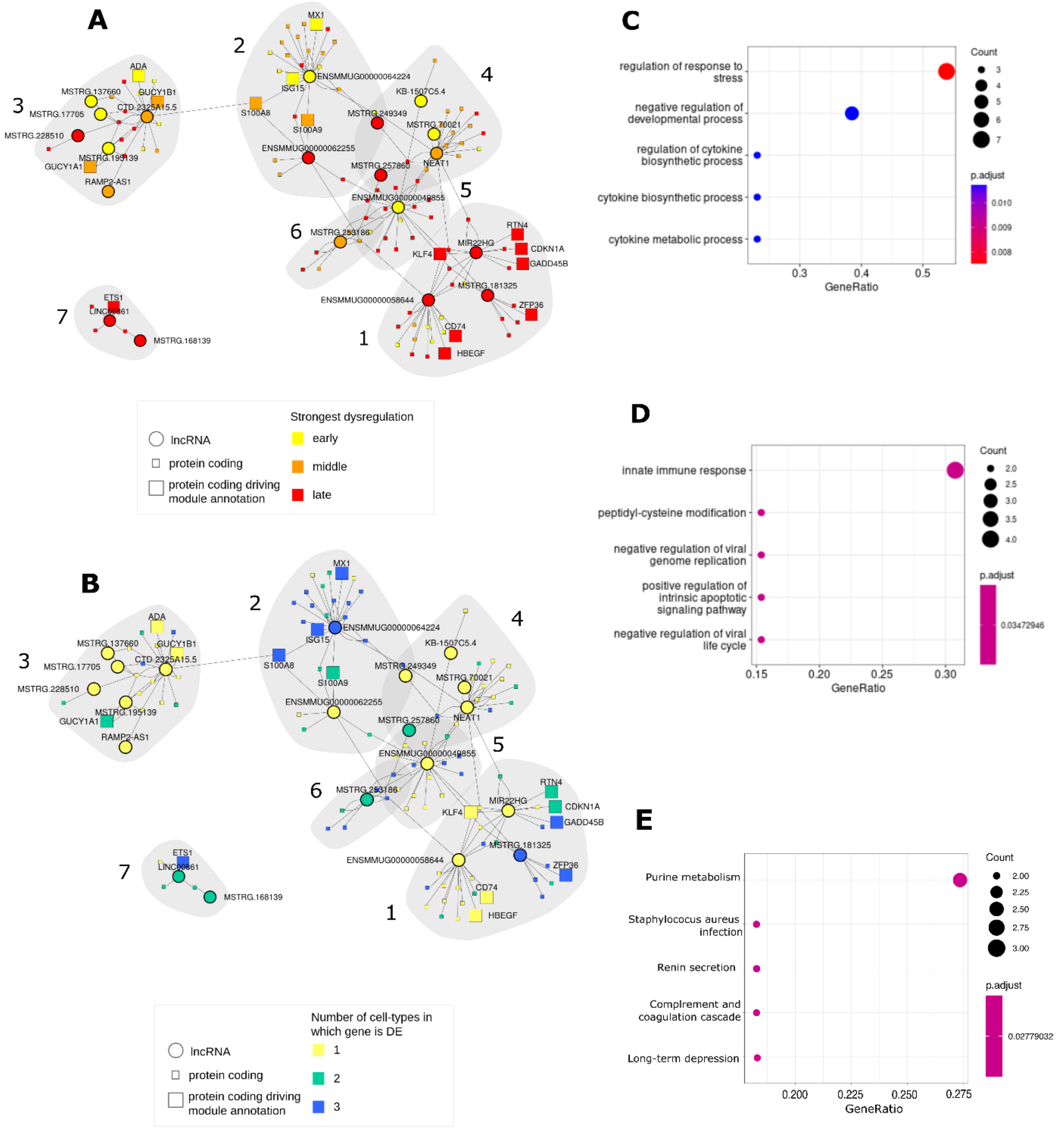


**Supplemental Figure S8. Co-expression network of lncRNAs and protein-coding genes upon EBOV infection in monocytes.** Regulatory network of DE lncRNAs (circles) and DE protein-coding genes (squares). **(A)** Vertices' colours represent whether a gene presents the strongest fold-change compared to baseline in early (yellow), middle (orange) or late (red) stage of infection **(B)** or whether they are DE in 1 (yellow), 2 (green) or 3 (blue) cell-types. GO functional enrichment of module 1 **(C)** and 2 **(D)**. KEGG functional enrichment of module 3. Protein-coding genes driving the functional enrichment of the module and lncRNAs are highlighted in bigger sizes.

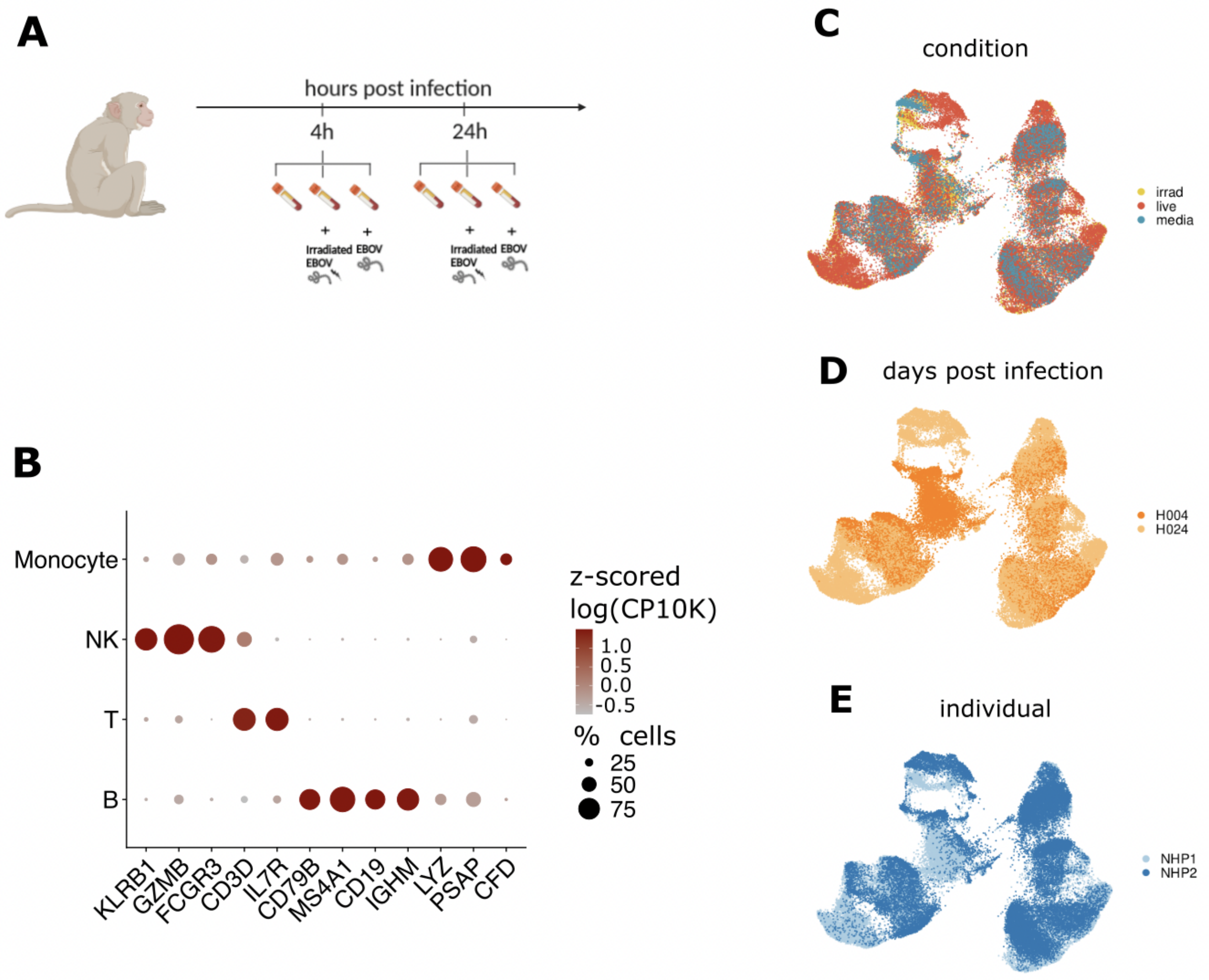


**Supplemental Figure S9.** ***Ex vivo*** **dataset processing.** **(A)** Schematic overview of the *ex vivo* experiment design. **(B)** Dot plot showing expression of cell-type markers which were used for cell type assignment in the *ex vivo* dataset. Dot's colours represent the average expression level of the gene in each cell type. Dot´s sizes represent the percentage of cells in which the gene is detected as expressed per cell type. UMAP embedding of cells from the *ex vivo* dataset coloured by condition (**C**), sampling hour relative to infection hour (**D**) and non-human primate (**E**).

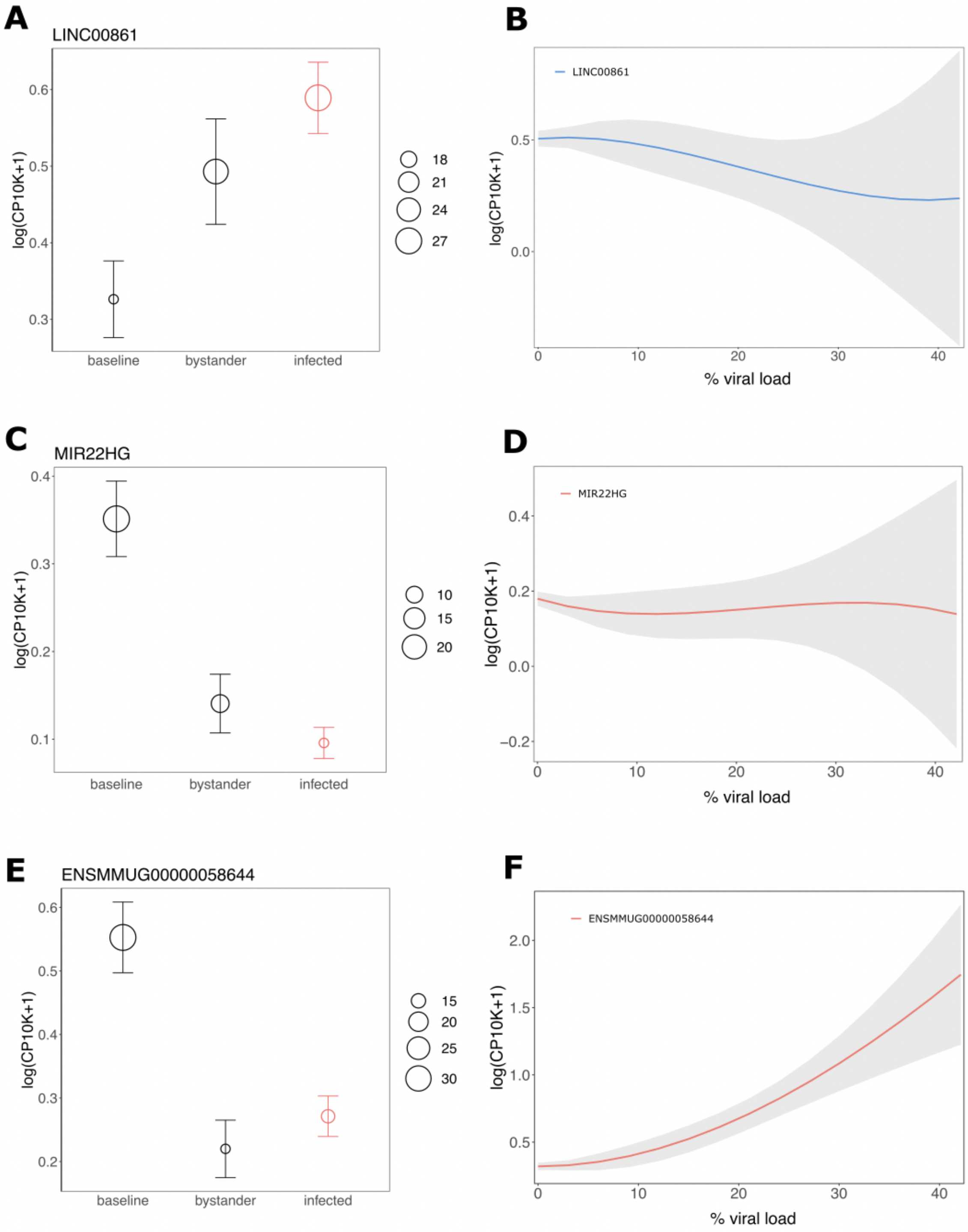


**Supplemental Figure S10. LncRNAs undergo expression changes within infected monocytes.**

Expression in baseline as well as bystander and infected cells at 24 hours post-infection. (Marker: mean; error bars: 95% confidence interval. Dot's sizes represent the percentage of cells in which the gene is expressed in each cell group) and expression changes with the viral load of LINC00861 **(A-B)**, MIR22HG **(C-D)** and ENSMMUG00000058644 **(E-F)**.
